# Analysis of stop-gain and frameshift variants in human innate immunity genes

**DOI:** 10.1101/003376

**Authors:** Antonio Rausell, Pejman Mohammadi, Paul J. McLaren, Ioannis Xenarios, Jacques Fellay, Amalio Telenti

## Abstract

Loss-of-function variants in innate immunity genes are associated with Mendelian disorders in the form of primary immunodeficiencies. Recent resequencing projects report that stop-gains and frameshifts are collectively prevalent in humans and could be responsible for some of the inter-individual variability in innate immune response. Current computational approaches evaluating loss-of-function in genes carrying these variants rely on gene-level characteristics such as evolutionary conservation and functional redundancy across the genome. However, innate immunity genes represent a particular case because they are more likely to be under positive selection and duplicated. To create a ranking of severity that would be applicable to the innate immunity genes we first evaluated 17764 stop-gain and 13915 frameshift variants from the NHLBI Exome Sequencing Project and 1000 Genomes Project. Sequence-based features such as loss of functional domains, isoform-specific truncation and non-sense mediated decay were found to correlate with variant allele frequency and validated with gene expression data. We integrated these features in a Bayesian classification scheme and benchmarked its use in predicting pathogenic variants against OMIM disease stop-gains and frameshifts. The classification scheme was applied in the assessment of 335 stop-gains and 236 frameshifts affecting 227 interferon-stimulated genes. The sequence-based score ranks variants in innate immunity genes according to their potential to cause disease, and complements existing gene-based pathogenicity scores.

**Author summary:** There are well characterized severe immunodeficiencies associated with loss-of-function variants in innate immunity genes. Genome sequencing projects identify rare stop-gain and frameshift variants in innate immunity genes whose phenotype is uncharacterized. Current methods to estimate the severity of rare stop-gains and frameshifts are based on evolutionary conservation of the gene and the likelihood for redundancy in its function. These parameters are not always applicable to innate immunity genes. We evaluated sequence-level characteristics of more than 30’000 stop-gains and frameshifts and prioritized variants according to their predicted functional consequences. Our scoring approach complements existing tools in the prediction of innate immunity OMIM disease variants and associates with functional readouts such as gene expression. In this framework, we show that many individuals do carry high pathogenicity variants in genes participating in antiviral defense. The clinical assessment of these variants is of significant interest.

## Introduction

There is considerable variability in the human immune response to pathogens. The observation of genetic causes of a number of primary immunodeficiencies underscores the fundamental role of variants in immune genes - in many cases resulting in severe, pathogen-specific disorders [1]. A main challenge in the analysis of genome variation today is the assignment of a functional role to rare variants. Here, large numbers of study participants would not necessarily provide the statistical power to associate a genotype with a phenotype. In this context, efforts are put toward to the computational identification of features allowing prioritizing variants for follow-up in genetic and functional analysis. Strategies to attribute a severity score to a variant, recently reviewed in [2], include approaches based on evolutionary, physico-chemical and structural properties ((Polyphen2 [3], SIFT [4]), methods based on analysis of mutation load (e.g. the Variation Intolerance Score [5]), and integrative pipelines [6-9].

Of special interest in the study of inter-individual variability in innate immunity is the evaluation of stop-gains and frameshifts. Such variants are prevalent, having an estimated number of 100 to 200 occurrences per human genome [10,11]. Stop-gains and frameshifts may lead to functional consequences due to protein truncation, degradation of the transcript by Nonsense-Mediated Decay (NMD) [12] and dominant negative influences of protein species. In particular, rare and young variants that have not undergone purifying selection may contribute to burden of disease in a population [13-15]. Despite a stop-gain or frameshift variant, however, the function of a protein may be preserved because of limited truncation of functional and structural domains, or because the variant affects only one of the splice forms. A less well understood possibility is the occurrence of stop-codon read-through [16,17].

Analyses based on gene characteristics such as evolutionary conservation and non-redundancy in the genome are generally used to predict the severity of stop-gain and frameshift variants ([18]). Herein, we will refer to these analyses as “*gene-based*”. However, innate immunity genes tend to be less conserved and more duplicated than the genome average [19] and other features may be needed to assess functional relevance of a variant. The aim of this study is to explore sequence characteristics that may improve the understanding of the functional consequences of stop-gain and frameshift variants in innate immunity genes. Herein, we will refer to these analyses as “*sequence-based*”. For this, we first evaluated two sets of publicly available data from a total of 7595 individuals [15,20] including gene expression data from 421 of them [21]. Specific sequence features of truncating variants were found to correlate with allele frequency and gene expression levels of affected individuals. These features were used to generate a pathogenicity score that was evaluated through benchmark against OMIM disease variants. The approach was applied to assess functional consequences of stop-gain and frameshift variants in innate immunity genes, with particular attention to antiviral interferon-stimulated genes (ISGs).

## Results

### Variant set

We analysed gene variant data from a total of 7595 individuals from the NHLBI GO Exome Sequencing Project (ESP) [15] and the 1000 Genomes Project [20]. We considered 17764 stop-gain and 13915 frameshift variants collectively affecting 11369 autosomal protein coding genes reliably annotated by the Consensus CDS (CCDS) project [22]. The distributions of gene truncating variants according to allele frequency and study are presented in **Supplementary Table S1**.

### Distribution of variants in sequence-based features

Consistent with previous reports [18,23], we observed that the distribution of stop-gain and frameshift variants along genes may be biased by allele frequency (**Supplementary Figure S1**). Variants with very low allele frequency (MAF≤0.001) are evenly distributed, with a modest 3’ terminal enrichment. However, the distribution of stop-gain and frameshift variants becomes less uniform with increasing allele frequencies, yet does not show a clear pattern. In contrast, we observed marked distribution trends in association with the following sequence features: (i) loss of functional domains; (ii) disruption of constitutive exons (i.e. exons present in all isoforms), or of principal isoforms; (iii) localization in potential NMD-targeted regions. In comparison to rare truncating variants, common stop-gain and frameshift variants were clearly depleted at positions leading to the loss of a functional domain (**Figure 1A**). Analysis of splicing-dependent effects was limited to genes with multiple annotated transcripts in CCDS (n=5203). We observed an enrichment of common stop-gain and frameshift variants in alternative isoforms (**Figure 1B**) and a depletion of common variants in principal isoforms (**Figure 1C**). We observed that common gene truncating variants occurred less frequently in regions more than fifty nucleotides upstream the last exon-exon junction, possibly triggering NMD-mediated transcript degradation (**Figure 1D**). For all the features discussed above, gene truncating variants associated with disease in the OMIM database exhibited a distribution bias opposite to what was observed for common stop-gain and frameshift variants (**Figure 1**).

**Fig 1.**
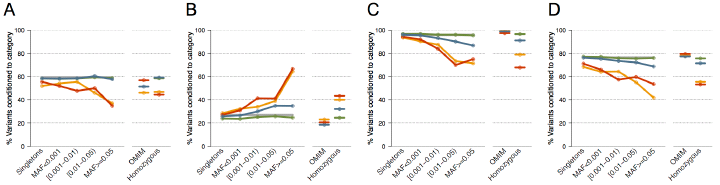
Distribution of variants according to sequence features and allele frequency: The percentage of variants upstream of a functional domain (**panel A**), in alternatively spliced sites (**panel B**), in the principal isoform (**panel C**) and in regions targeted by NMD (**panel D**). The distribution is shown for synonymous (green), missense (blue), stop-gains (red) and frameshift (orange) variants according to minor allele frequency (MAF) intervals, where singletons (variants detected only in one individual) are represented separately. The pattern of OMIM disease variants and homozygous variants for each feature is shown. The corresponding coding genome background (measured as the percentage of nucleotides displaying the feature) is shown as a grey line (partly hidden by the distribution of synonymous variants in some panels). Numbers of variants in each category are reported in **Supplementary Table S2**. Logistic regression was used to model the relationship between observing a given sequence feature in a given type of variant as a function of the logarithm of the minor allele frequency (MAF). The odds ratio estimates for stop-gain variants were significantly different from those of synonymous variants in all panels (p-values<5e-05, heterogeneity test [36]; for frameshifts, in panels B, C and D (p-values<5e-03).

### Expression analysis

We used expression data from 421 individuals to assess the functional impact of stop-gain and frameshift variants [21]. In particular, we evaluated differences between protein truncation variants localized or not in the NMD-targeted region. Stop-gains predicted to trigger NMD (n=756) had a median Z-score =−0.59, significantly lower than stop-gains predicted to escape NMD (n=379, median=−0.10) and lower than a reference distribution of synonymous variants (median =−0.04, one-sided Wilcoxon rank-sum test p-value < 2.2e-16) (**Figure 2A**). Among stop-gains predicted to trigger NMD, singletons (n=488, median Z-score=−0.75) showed a higher decreased in gene expression levels than non-singletons (n=268, median=−0.26, p-value=1.3e-10, **Figure 2B**), which is an indication that they represent actual variants and not sequencing or bioinformatics errors. We did not observe a similar reduction when considering 87 of the 172 frameshift variants with expression data mapping to potential NMD-target regions (median=−0.14, p-value=5.7e-02).

**Fig 2.**
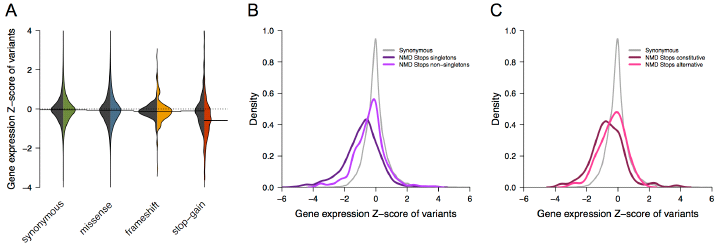
Association of NMD-target variants with gene expression. **Panel A** shows the distribution of average expression z-scores for genes from individuals carrying different types of variants (synonymous, missense, frameshift and stop-gain). Peer-factor normalized RPKM from [21] were used. The black half represents the distribution of variants outside the NMD-target region and the colored half for those within the NMD-target region. Statistically significant differences were observed for stop-gain variants predicted to trigger NMD (n=756) compared to synonymous variants (one-sided Wilcoxon rank-sum test p-value < 2.2e-16). **Panel B** shows the distribution of average expression z-scores described in panel A for synonymous (grey) and stop-gain (dark and light purple) variants within the NMD-target region. The distribution of NMDtarget stop-gains is represented separately for singletons (dark purple, n=488) and nonsingletons (n=268). Distributions are statistically different (one-sided Wilcoxon rank-sum test =1.3e-10). **Panel C** shows the distribution of average expression z-scores described in panel A for synonymous (grey) and stop-gain (dark and light pink) variants within the NMD-target region of genes with multiple isoforms described in CCDS. The distribution of NMD-target stop-gain is represented separately for those affecting all isoforms (dark pink, n=216) and those affecting only a fraction of isoforms (light pink, n=85). Distributions are statistically different (one-sided Wilcoxon rank-sum test = 2.5e-03). Results were reproduced using RPKM normalized expression values (**Supplementary Figure S2**).

To further evaluate a splicing-dependent impact in gene expression levels, we limited the analysis to 301 stop-gains predicted to trigger NMD and affecting genes with multiple isoforms described in CCDS. We observed a significant decrease in gene expression levels of NMD-triggering stop-gains affecting all isoforms (n=216, median=−0.64) compared to those affecting only a fraction of isoforms (n=85, median=−0.22, one-sided Wilcoxon rank-sum test p-value = 2.5e-03) (**Figure 2C**). Similar results were obtained using RPKM normalized expression values (**Supplementary Figure S2**). These observations confirmed the functional impact of stop-gains consistently with predictions of degradation by NMD and current annotation of isoforms.

### Pathogenicity scores

We then evaluated the predictive value for pathogenicity of the sequence-based features characterized in previous sections: percentage of sequence affected, loss of functional domains, proportion of isoforms affected, principal isoform damage, and NMD-target region. We integrated them into a naïve Bayes classifier and assessed its performance over a dataset of 1160 pathogenic stop-gain variants found in OMIM database and 125 common stop-gain variants that are not known to be pathogenic. Predictive performance of the pathogenicity score was validated over unseen variants excluded from training data using successive random subsampling (see Methods). The classifier was benchmarked against a state of the art gene-based probability score proposed by [18]. This gene-based score relies on conservation and protein interaction network proximity to genes associated to a recessive disease as predictive features. In the case of stop-gain variants, the performance of the gene-based method was consistent with the reported results in the original work (Area Under the Curve (AUC) = 0.83, **Figure 3**). The score based on sequence features alone also showed predictive value (AUC = 0.67). Additionally, we tested if a combination of the two scores resulted in an improvement of the classification power. Indeed, the optimal ROC was achieved by combining the two scores (AUC = 0.87). Similar results were obtained for frameshift variants (**Supplementary Figure S3**). These results demonstrate that sequence features can be incorporated as an additional source of information to improve current pathogenicity prediction.

**Fig 3.**
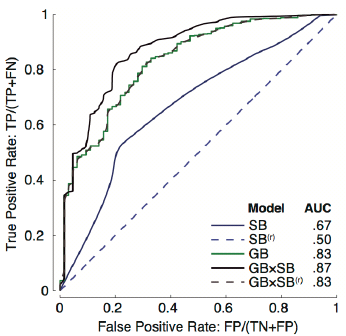
Receiver operating characteristic of the performance of pathogenicity scores for stop-gain variants. Classification power of three pathogenicity scores was evaluated on a set of 1160 pathogenic stop-gain variants found in the OMIM database, and 125 common stop-gain variants not known to be pathogenic. Shown are the ROC curves for the sequence-based classifier (SB) developed in this work, for the gene-based score reported in [18] (GB), and for the joi t classifier (SB×GB). Dashed curves correspond to a randomization test in which rows in sequence features are shuffled column-wise (denoted by SB^(r)^). Incorporating sequence features leads to an increased area under the ROC curve. A similar plot using frameshift variants is given in **Supplementary Figure S3**.

### Analysis of innate immunity genes

To test the ability to rank the functional consequences of gene truncating variants in innate immunity genes, we analysed the distribution of both the sequenced-based and gene-based pathogenicity scores in 1503 genes involved in innate immunity [19], including 387 interferon stimulated genes (ISGs, [24] [25]). We identified 856 innate immunity genes, including 230 ISGs, carrying rare gene truncating variants (MAF<1%). Globally, innate immunity and OMIM genes ranked higher than the background set of the genome for both scores (**Figure 4** **and Supplementary Figure S4**). However, the highest scores were obtained for stop-gain variants in OMIM genes, particularly for variants in innate immunity genes that are not observed in the ESP or the 1000 Genomes Project samples (**Figure 4**). The latter result is consistent with their extreme rarity and severity. We note that OMIM variants used here for validation were not considered for learning in the Bayesian classification (see Methods). Despite their apparent agreement in **Figure 4**, correlation between the sequenced-based and gene-based pathogenicity scores was very low (Spearman correlation coefficient below 0.31 in all sets of genes analyzed) indicating that both scores provide complementary information.

**Fig 4.**
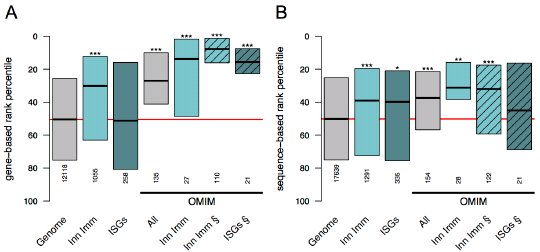
Pathogenicity score distributions for rare stop-gain variants in innate immunity genes. Rank percentile distributions of pathogenicity scores for rare stop-gain variants (MAF<1%) are shown in different sets of genes: protein coding genome background (grey, “Genome”), innate immunity genes (light turquoise, “Inn Imm”) and their subset of interferon stimulated genes (dark turquoise, “ISGs”). The same categories are shown for OMIM disease variants. All variants are reported in ESP and 1000 Genomes Projects except for sets indicated with the § symbol (dashed boxes) which present scores for OMIM disease variants only reported in the OMIM database. Only three variants reported in ESP and 1000G were found to affect ISGs and be annotated as pathogenic in OMIM. Therefore this last category is not represented in the figure. Variants with the highest probability of being pathogenic have rank percentiles closer to zero (top of the panels). **Panel A** represents precomputed gene-based pathogenicity scores from [18]. **Panel B** represents sequence-based pathogenicity scores, i.e. posterior probabilities using the features described in the present work (see main text). Each box spans between 1st and 3rd quantile, and the median is denoted by a bold line in the middle. Total number of variants within each distribution is indicated. Differences in number of variants in equivalent categories between panel A and B originate from unavailability of the gene-based scores for some genes. Statistical differences against the genome reference (one-sided Wilcoxon rank sum tests) are indicated with asterisks according to Bonferroni corrected pvalues: <5e-02 (*), <5e-03 (**) and <5e-04 (***). The genome-wide median is denoted by a red line. Spearman correlation coefficient between the sequenced-based and gene-based pathogenicity scores was below 0.31 in all sets of genes analyzed.

We then focused on gene truncating variants in genes associated with viral inhibition in cellular assays [24,25]. A total of 13 out of 42 genes carried such variants (observed in at least 2 people), which were very rare overall (MAF<0.0053; **Table S3**). Specifically, two genes had variants with high predicted pathogenicity based on both scores: *MX1* which controls Influenza A virus *in vitro* and *HPSE* which is involved in metapneumovirus, respiratory syncytial virus and yellow fever virus control. While the gene-based scores were by definition identical for all variants affecting a same gene, the sequenced-based score sharply distinguished the variants according to different predictive pathogenicity (**Table S3**). This observation was consistent with the observed differences in gene expression levels available for some of the variants.

## Discussion

Numerous Mendelian disorders leading to severe infection are caused by rare functional variation of innate immunity genes [1]. Here, we identified multiple stop-gain and frameshift variants in this family of genes in the general population, especially among interferon stimulated genes. These are generally heterozygous rare variants that may or may not result in clinical consequences. To understand the nature and possible consequences of these variants, we first analyzed their characteristics at the genome level. The genome-wide analysis of more than 30’000 variants provided the statistical power to identify sequence specific features for severity and to build a pathogenicity score. This sequence-based pathogenicity score was then applied to the analysis of variants in interferon stimulated genes with antiviral activity.

We observed that the distribution of stop-gain and frameshift variants in the sequence is biased by the allele frequency. Thus, we speculated that tolerance to these variants would reflect their impact on functional domains, on isoforms, and on degradation by NMD. Our results clearly underscore that rare stop-gain and frameshift variants are subject to purifying selection [14,26]. Indeed, those variants are kept at very low frequency when they result in the loss of functional domains, when they are located in NMD-targeted regions, or when they disrupt the principal isoform or constitutively spliced exons. The potential molecular impact of heterozygous rare truncating variants was examined using mRNA expression data [21]. Stop-gain variants predicted to trigger NMD degradation resulted in a measurable decrease in global expression levels. This is in line with recent findings showing a reduction in expression levels of the variant allele compared to the reference allele in heterozygous individuals when stop-gains occur in NMD target regions [18,21,27]. In all analyses, singleton variants associated with highest functional impact, consistent with higher severity of lower frequency rare variants and indicative of general accuracy in variant calling.

To further explore the functional consequences of gene truncating variants, we analysed the collective contribution of various severity features to the prediction of pathogenicity. For this we built a model on a training set that was validated through benchmark against OMIM disease variants. These sequence-based features improved the ranking of OMIM variants when added to a predictive model that use gene-based features. We hypothesized that such sequence-based approach would be of particular interest for the study of innate immunity genes because, as a group, these genes tend to be less conserved than the genome average and hence need special consideration. The analysis showed that our sequence-based score is able to rank variants in innate immunity genes according to their pathogenicity and provides complementary information to previously proposed gene-based scores. Indeed we found that in the case of the antiviral genes *MX1* and *HPSE*, truncating variants ranked very high in pathogenicity on the basis of gene-based scores while important differences were observed at sequence level suggesting significant differences in functional impact. For example the *MX1* stop-gain rs35132725 exhibits all the features of severity and a negative effect on expression levels. In contrast, the *MX1* frameshift rs199916659 is not expected to alter protein function.

Overall, among 387 ISGs examined in 7595 individuals, more than half of the genes carried a stop-gain or frameshift variant in 1 or more individuals, usually at low allele frequency. We then evaluated those instances that concerned genes for which an antiviral effect has been established through a gain-of-function screen *in vitro*. This last analysis provided a short list of genes and reliable variants that could modulate responses to various viruses, including common human pathogens such as influenza. Of note, the *in vitro* virological inhibition data represents a technical readout, and there are a number of considerations that may diminish the *in vivo* consequences of these rare variants, including issues of redundancy and robustness in innate immunity networks, and the possibility of stop codon read-through. There are other limitations to the predictions based on sequence features, particularly the incomplete understanding of the functional role of alternative isoforms and their tissue specificity.

Rare gene truncating variants predicted to have high pathogenicity risk in innate immunity genes should be examined for phenotypic consequences in the population. Exceptional homozygous individuals may be at risk for severe infection while heterozygous individuals could have adequate compensation or subtler phenotypes. However, there is increasing awareness of the relevance of haploinsufficiency [28], and thus, it is not excluded that heterozygosity may be associated with apparent clinical phenotypes. Thus, the next step should include assessment *in vivo* of high risk variants, which requires the capacity to re-contact carrier individuals for collection of biological specimens and in-depth phenotypic assessment.

## Material and Methods

### Human variation sets

Two genetic variant and annotation datasets were used: 1) 6503 individuals from the NHLBI GO Exome Sequencing Project (ESP) [15] and 2) 1092 individuals from the 1000 Genomes Project [20]. Variants (SNPs and INDELs) and annotations for the ESP exomes (file ESP6500SI-V2-SSA137.dbSNP138-rsIDs.snps_indels.txt.tar.gz) were downloaded from the Exome Variant Server, NHLBI GO Exome Sequencing Project, Seattle, WA (http://evs.gs.washington.edu/EVS/, accessed July 2013). Only variants assigned to the following categories were considered for further analysis: “stop-gained” (including “stop-gained-near-splice”), “frameshift”, “coding-synonymous” (including “coding-synonymous-near-splice”) and “missense” (including “missense-near-splice”). One base was added to the genomic coordinates reported for frameshifts in the ESP dataset to consider the actual location of the insertion/deletion event (http://evs.gs.washington.edu/EVS/HelpDescriptions.jsp?tab=tabs-1). Variants and genotypes from the 1000 Genome Project [20] correspond to phase 1 version 3 of the 20110521 release (ftp://ftp.1000genomes.ebi.ac.uk/vol1/ftp/release/20110521/, accessed August 2013). SnpEff Variant Analysis software [29] (version 3.3h build 2013-08-11) was used to annotated 1000Genome variants against SnpEff’s pre-built human database (GRCh37.71). SnpEff categories labeled with errors or warnings in the EFF field were disregarded. Only variants assigned to the following categories were considered for further analysis: “stop_gained”, “frame_shift”, “synonymous_coding” (including “synonymous_start” and “synonymous_stop”) and “non_synonymous_coding” (missense). Hardy-Weinberg equilibrium (HWE) was tested with R package GWASExactHW (http://cran.r-project.org/web/packages/GWASExactHW/, version 1.1). A fraction of variants significantly deviated from HWE (Fisher’s exact p-values <0.05), mainly due to an excess of homozygous rare allele calls, likely indicating technical artifacts. All variants not in HWE were filtered out. When both datasets where considered together, the following criteria were adopted: i) Genomic coordinates of frameshift variants reported by both datasets were treated as reported for the ESP dataset. ii) Allele frequencies and HWE of variants present in both datasets were derived from the sum of individuals from both studies; allele frequencies of variants present in only one dataset were taken as originally reported by the corresponding dataset. To exclude bias due to previous assumptions, results were reproduced for the two datasets considered separately as well as combined considering the allele frequencies of variants present in only one dataset are estimated by including 7593 individuals as the common denominator.

### Annotation of variants in reference human transcript and protein sequences

The analysis pipeline implemented to annotate genetic variants is depicted in **Supplementary Figure S5**. We restricted the analysis to protein coding genes and transcripts annotated by the Consensus CDS (CCDS) project [22] (ftp.ncbi.nlm.nih.gov/pub/CCDS/, Release 12 04/30/2013). We considered only variants affecting a core set of human protein coding regions consistently annotated and of high quality. Only genes on the 22 autosomal chromosomes were retained and only CCDS entries with a public status and an identical match were kept.

Domains of human protein sequences were retrieved from the InterPro database [30] (release 44.0, 23/09/2013). Data were downloaded through BioMart Central Portal [31] (http://central.biomart.org/, accessed 04/10/2013), filtering fragments and considered domain boundaries corresponding to InterPro “supermatches”. Mapping from InterPro coordinates on UniProt protein sequences to CCDS sequences was done by exact matching of the complete amino acid sequences using UniProt database (release 2013_07; [32]).

A position within a protein coding gene was considered alternatively spliced if it was shared by only a fraction of all protein coding transcripts reported by the Consensus CCDS Project for that gene. Otherwise it was considered constitutively spliced for the purpose of the study. Annotation of principal isoforms used APPRIS ([33]; file APPRIS-g15.v3.15Jul2013/appris_data.principal.homo_sapiens.tsv accessed 03/09/2013 at URL: http://appris.bioinfo.cnio.es), a computational pipeline and database for annotations of human splice isoforms. APPRIS selects a specific transcript as principal isoform, i.e. the one computationally predicted as responsible of the main cellular function, being expressed in most of the tissues or developmental stages and more evolutionary conserved. Selection of the principal isoform is based on protein structure, function and interspecies conservation of transcripts.

We accounted for nonsense-mediated decay (NMD) following HAVANA annotation guidelines v.20 (05/04/2012) (http://www.sanger.ac.uk/research/proiects/vertebrategenome/havana/assets/guidelines.pdf), Specifically, the NMD-target region of a transcript was defined as those positions more than 50 nucleotides upstream the 3’-most exon-exon junction. Transcripts bearing stop-gain variants at these regions are predicted to be degraded by NMD [12].

### Functional validation using mRNA expression data

Geuvadis RNA sequencing data from 421 lymphoblastoid cell lines from the 1000 Genomes Project (phase 1 version 3 of the 20110521 release, see above; [20]) were obtained from Lappalainen et al. 2013 [21]. Gene and transcript expression quantifications of protein-coding genes were downloaded from EBI ArrayExpress accession E-GEUV-1 (accessed 05/11/2013). Analyses were independently performed on both RPKM and Peer-factor normalized RPKM values. As a measure of the impact of a variant on expression level, we calculated the average Z-score of the expression level in cells from individuals carrying the variant compared to all samples.

### Derivation of sequence-based pathogenicity score

We used a naïve Bayes classification scheme in order to derive a probability of pathogenicity for a given variant using the following sequence-based features: maximum transcript length affected, maximum percentage of domain truncation, number of isoforms and ratio of isoforms affected, truncation of the principal isoform and localization in an NMD-target region. Solely for the purpose of the classifier, missing values were imputed to zero for percentage of domain truncation and to the longest isoform for principal isoform annotation. We defined a matrix *X*_N×K_ of *K* sequence-based features for *N* variants of a given type in the dataset and a binary vector ***c***_N×1_ annotating variants as benign or pathogenic. A new variant, ***y***_1×K_, is evaluated using maximum likelihood estimates for class-specific means from the annotated data, and a common intra-class variance vector (except for binary features). We estimate the variance vector as, 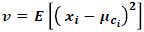, where ***x***_*i*_ is the *i*^*th*^ row in matrix *X* and 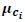 is the mean vector corresponding to the class indicated by *c*_*i*,_. We assigned a pathogenic class with 1 and benign class with 0. Assuming a prior probability of pathogenicity, *p*_*1*_, posterior probability of pathogenicity can be evaluated as, 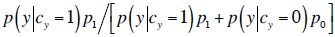, where ***p***_**0**_ = **1** – ***p***_**1**_ is the prior probability of being benign. The conditional likelihood of ***y*** for a given class is assumed to factorize as product of *K* likelihoods corresponding to the *K* sequence features available (naïve Bayes assumption). We used normal, and Bernoulli likelihood functions to model continuous and binary features respectively. It is straightforward to show the ranking produced from this posterior probability does not depend on the prior probability *p*_*1*_ as long as it is larger than zero and it is equal for all the mutations under consideration.

### Evaluation of pathogenicity scores

As reference throughout the work, and as a training set for the predictive scores, we used a catalogue of pathogenic mutations from the Online Mendelian Inheritance in Man [34] database. Only genes with a cytogenetic location (genemap2.txt accessed 18/10/2013 at OMIM: ftp.omim.org) and with a gene status of confirmed or provisional were kept. For each gene with an associated MIM number, all allelic variants with a “live” status and a dbSNP identifier were obtained through the OMIM API server (http://api.omim.org/). We used Ensembl Variation [35] (Ensembl release 71, April 2013, dataset Homo sapiens Short Variation, SNPs and indels, GRCh37.p10, accessed 25/10/2013 at http://apr2013.archive.ensembl.org/biomart/martview/) to obtain the genomic coordinates for each dbSNP identifier together with the clinical significance of each specific allele as reported by ClinVar and dbSNP following OMIM guidelines (http://www.ncbi.nlm.nih.gov/clinvar/docs/clinsig/). Only variants with a dbSNP identifier annotated as “pathogenic” and mapping to a unique genomic location were kept for further analysis. SnpEff Variant Analysis was then used to re-annotate the selected pathogenic variants as described above.

We benchmarked three different pathogenicity scores using all stop-gain variants from the OMIM dataset as positive (pathogenic), and all common variants (MAF ≥ 1%) not present in OMIM dataset as negative (benign) variants. The sequence-based score is the posterior probability calculated from the naïve Bayes classification scheme described in previous section using an empirical prior for pathogenicity. The gene-based score is the probability provided by MacArthur et al [18] for prioritization of variants derived from two gene-level features: conservation and protein interaction network proximity to genes associated with a recessive disease. The joint score is defined as the product of the two previously defined probability scores. This joint probability is the probability of a pathogenic mutation in a disease gene assuming conditional independence of the two probability scores. The receiver operating characteristic (ROC) curve was derived using random subsampling validation iterations. In each iteration, we use 75% of the data to train the classifier and use the remaining 25% for validation. This was done 10000 times, to minimize the Monte Carlo error, and validation set scores were combined to calculate the ROC curve. The same procedure was applied to frameshift variants.

### Analysis of innate immunity genes and interferon stimulated genes (ISGs) with antiviral activity

A representative list of 1503 human innate immunity genes [19] was used. Within this list, we further analyzed 387 interferon stimulated genes (ISGs) [24]. Additionally, we focused on those ISGs showing antiviral activity against 18 viruses (including important human pathogens such as HIV-1, hepatitis C virus, influenza virus and other respiratory viruses) upon overexpression in *in vitro* cellular assays [24,25]. We first identified all ISGs carrying gene truncating variants, and then characterized the subset of those genes associated with more than 50% viral inhibition in the cellular assays.

## Acknowledgments

The authors would like to thank Zoltan Kutalik for feedback on statistical analyses, the 1000 Genomes Project, the Geuvadis Consortium and the NHLBI GO Exome Sequencing Project. Some of the computations for this study were performed at the Vital-IT (http://www.vital-it.ch) center for high-performance computing of the Swiss Institute of Bioinformatics. All perl, Matlab and R scripts developed for this work are available upon request.

